# Fast Hierarchical Bayesian Analysis of Population Structure

**DOI:** 10.1101/454355

**Authors:** Gerry Tonkin-Hill, John A. Lees, Stephen D. Bentley, Simon D.W. Frost, Jukka Corander

## Abstract

We present fastbaps, a fast solution to the genetic clustering problem. Fastbaps rapidly identifies an approximate fit to a Dirichlet Process Mixture model (DPM) for clustering multilocus genotype data. Our efficient model-based clustering approach is able to cluster datasets 10-100 times larger than the existing model-based methods, which we demonstrate by analysing an alignment of over 110,000 sequences of HIV-1 pol genes. We also provide a method for rapidly partitioning an existing hierarchy in order to maximise the DPM model marginal likelihood, allowing us to split phylogenetic trees into clades and subclades using a population genomic model. Extensive tests on simulated data as well as a diverse set of real bacterial and viral datasets show that fastbaps provides comparable or improved solutions to previous model-based methods, while generally being significantly faster. The method is made freely available under an open source MIT licence as an easy to use R package at https://github.com/gtonkinhill/fastbaps.

## Introduction

Identifying clusters of genetically similar individuals within a larger population is a common problem in genetics and ecology. Population structure is helpful in understanding past historical population events, conservation genetics, the analysis of invasive species and disease outbreaks. Confounding population structure must also be considered in tests for natural selection as well as genetic association studies (1–3).

Methods for clustering genotype data can generally be separated into those based on a statistical model of population structure, where observations from each underlying cluster are assumed to be drawn from a parametric model, and distance-based methods that rely on more general clustering techniques such as k-means (4). Whilst distance-based approaches are generally faster, they are not as readily interpretable and come with no statistical guarantees regarding large-sample behavior. Model-based methods typically indicate the probability that an individual belongs to a certain cluster, allow for different models to be compared and can provide estimates of uncertainty for the inferred parameters. Generally, either Bayesian or maximum-likelihood based approaches are widely used for inference in population structure models.

A commonly considered model of population structure assumes that there is a fixed number of uncorrelated underlying populations, *K*. An individual is then assumed to originate either from a single population, the no-admixture model, or to carry alleles from multiple populations, which corresponds to the admixture model (5). Solutions to this problem include STRUCTURE (5, 6), BAPS (7–9), Admixture (10), fastStructure (11) and the method of Anderson and Thompson (12). More recently, methods that combine model-based techniques with an initial faster distance-based clustering have been proposed including snapclust (13) and hierBAPS (14).

A common theme of most model-based methods is that they require the underlying number of clusters, *K*, to be provided. In practice, the methods are usually run over a range of *K* and the ‘best fitting’ model is selected (15, 16). An alternative Bayesian solution is to put a prior on the number of clusters, usually using a Dirichlet process mixture model (DPM) in an attempt to infer *K* as part of the model (17, 18). Whilst this provides a natural method for inferring an appropriate *K*, the approach is very computationally expensive and does not scale to large numbers of loci and individuals. The greedy stochastic optimisation approach in hierBAPS (14) places a discrete uniform prior on *K* up to a supplied maximum value and is able to scale to larger datasets. However, as datasets become very large, inferring *K* using model-based clustering approaches has so far been infeasible. An example of a dataset that is too large for current model-based methods is the set of HIV-1 pol genes, which are routinely generated in clinical settings for the purposes of identifying resistance to antivirals. Despite the relatively short length of this region (typically 1200-1500 base pairs long), the very large number of sequences - in excess of 100,000 in public databases alone-makes choosing an appropriate value for *K* very challenging.

The Bayesian hierarchical clustering (BHC) algorithm (19) presents an alternative approach that approximates a Dirichlet Process Mixture model whilst guaranteeing a solution in polynomial time. The method performs a version of agglomerative bottom up clustering using a Dirichlet process to account for uncertainty in the data and Bayesian model selection to decide which clusters should be merged at each step. Whilst the approach has been shown to succeed in clustering text documents, microarray data and even electrical demand profiles (19–21), it has yet to be successfully applied to the problem of identifying population structure. A reason for this is that the hyperparameters (*β*) of the model defining each component of the mixture are set to be proportional to the counts of each allele at each loci in the complete dataset. In practice setting the hyperparameters to be proportional to the entire dataset leads to an overpartitioning of population genetic data whereby the algorithm identifies a very high number of clusters (see Results). A common alternative prior in population structure studies is to use a symmetric Dirichlet prior (5). However, at the lowest levels of the hierarchy a symmetric prior will lead to different clustering combinations having the same posterior likelihood. This greatly reduces the ability of the algorithm to accurately cluster sequences at the lower levels. In contrast, the hierBAPS algorithm relies on a fast initial clustering using complete-linkage agglomerative clustering to provide an initial partition of the dataset. hierBAPS then performs a greedy stochastic optimisation procedure to identify the local maximum a posteriori (MAP). By taking advantage of a fast initial clustering approach similar to that used in hierBAPS we are able to place a symmetric or BAPS prior on the mixture components, enabling the BHC algorithm to distinguish between different combinations of clusters.

Though methods such as hierBAPS and snapclust have allowed for analyses to scale to much larger datasets than previous approaches, they remain in practice inapplicable to the currently emerging datasets comprising tens of thousands or even hundreds of thousands of sequences, where very large numbers of underlying clusters may be present. Here, through incorporating ideas from both BHC and hierBAPS, we produce an efficient inference solution to the no-admixture model for very large datasets when the underlying number of clusters may be in the tens or hundreds.

## Methods

### Initial clustering

Similar to hierBAPS, but unlike the original BHC algorithm, fastbaps begins by generating a fast initial clustering of the data using a value for *K* much larger than expected (*K*_init_). This has the advantage of both reducing the complexity of the Bayesian agglomerative stage whilst allowing for symmetric priors to be used. Starting with a multiple sequence alignment in FASTA format, we first generate a pairwise single nucleotide polymorphism (SNP) distance matrix before clustering using Ward’s agglomerative clustering (22). This hierarchy is cut to generate *K*_init_ clusters. The hierarchy is also used to estimate the hyper-parameters *β* of the individual mixture component priors. Very large datasets make the calculation of the initial pairwise distance matrix prohibitive. To counter this limitation, for datasets with more than 10,000 individuals we produce the initial clustering by first performing a Principal Component Analysis before clustering the first 50 principal components using a fast hierarchical algorithm such as genie or the memory efficient Ward algorithm in the fastcluster package (23, 24). This removes the need to calculate the complete distance matrix.

### Bayesian hierarchical clustering

Given a collection of small clusters and their respective hierarchies, the BHC algorithm of Heller and Ghahramani (19) proceeds in a similar fashion to traditional agglomerative clustering. In the place of a distance metric, Bayesian hypothesis testing is conducted at each level to decide which clusters to merge. Let 
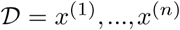
 describe the entire dataset of *n* samples and 
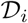
 the set of points at the leaves of the subtree *T_i_*. We initialise the algorithm with the small subtrees obtained using the fast clustering step as shown in Figure 1. At each subsequent stage we then consider merging all pairs of existing trees. If trees *T_i_* and *T_j_* are merged into a combined tree *T_k_* then the set of points corresponding to the new tree is 
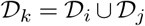
. At each merge we compare two hypotheses. The merged hypothesis, which we denote 
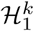
, is that all data in 
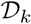
 were generated identically and independently from the same probabilistic model *p*(***x***|*θ*) with unknown parameters *θ*. In our case this model is a multinomial, with parameters *θ* = (*n, p*). We also specify a Dirichlet prior over the parameters *p* with hyperparameters *β*. Thus, we can write the probability of the data 
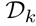
 under 
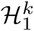
 as

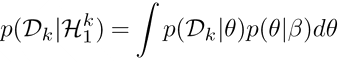

**Fig. 1.**
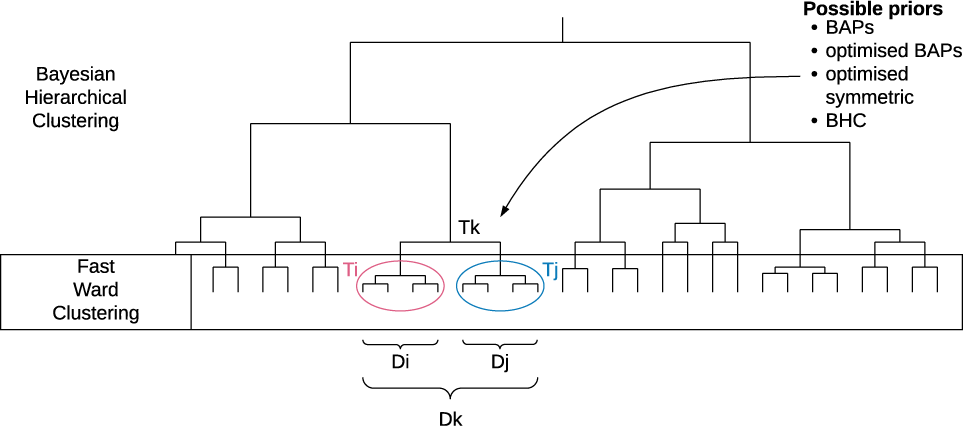
A diagram outlining the fastbaps algorithm. An initial clustering is performed by default using agglomerative clustering with the Ward linkage method. Subsequently the merge with the highest posterior probability is chosen. The final clustering is then determined by only accepting merges with a posterior probability > 0.5

Assuming a multinomial-Dirichlet distribution, this integral is tractable and we can rewrite the above equation as

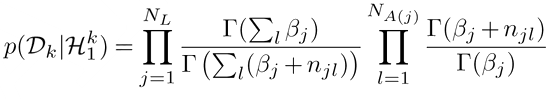

Where *N_L_* is the number of loci, *N_A_(*j*)__* is the number of possible alleles at loci *j* and *n_jl_* is the number of copies of allele *l* at locus *j*. *β_j_* is the corresponding Dirichlet prior hyperparameter.

The alternative hypothesis to merging two clusters 
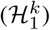
 is that the data 
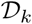
 have two or more clusters. Summing over all the possible partitions of 
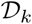
 into two or more groups is in practice intractable. However, by only considering partitions that are consistent with the subtrees *T_i_* and *T_j_*, we can use recursion to quickly compute this sum. Under the restriction of remaining consistent with the subtrees, the probability of the data under the alternative hypothesis 
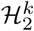
, is just the product over the subtrees 
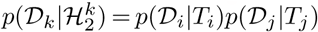
 (see Figure 1) where the probability of the data given the tree is given by

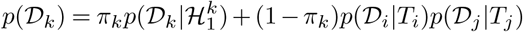

The prior for the merged hypothesis 
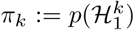
, can be computed bottom-up as described in Heller and Ghahramani, (19) and is a deterministic function of the concentration parameter of the DPM. In practice, for the size of datasets we are considering, the actual value of *α* makes very little difference as the prior *π_k_* is dominated by the factorial on the number of points within a cluster. Similar to Savage et al., (21), we fix t he v alue o f *α*, the concentration parameter of the DPM, rather than keeping it as a user-tunable hyperparameter in the resulting clustering algorithm.

Finally, given the probabilities of each hypothesis described above we can calculate the posterior probability of the merged hypothesis 
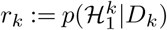
 using Bayes’ rule

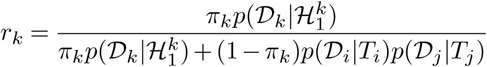

We use this quantity to greedily decide which two subtrees to merge. After generating a final hierarchy we then use the same quantity to identify which merges were justified and cut the tree when *r_k_* < 0.5.

### Hyperparameter selection

As described in Heller and Ghahramani (19), for any given set of hyperparameters, the root node of the hierarchy approximates the probability of the data given that setting of the hyperparameters. By leveraging this, we use the marginal likelihood of the root node 
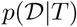
 to perform model comparisons for different settings of the hyperparameters. We found that different settings of the hyperparameter *α* had a negligible impact on the final clustering due to the dominance of the factorial component of *π_k_*, which provided a strong motivation to fix the hyperparameter at the value *α* = 1. Similar to Heller and Ghahramani (19) and Savage et al., (21), we use golden section search as provided in R’s ‘optimise’ function to select the *β* parameters and thus the variance of the Dirichlet prior. Unlike previous approaches, we did not set the *β* parameters to be proportional to the discrete total counts for the entire dataset. Instead we used either a symmetric prior, similar to Pritchard et al.,(5) or the non-informative prior 
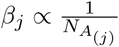
 as used in hierBAPS and suggested in Anderson and Thompson (12). These priors are referred to as optimised symmetric, and optimised BAPS priors hereafter. These priors were found to generally outperform the prior used in the original BHC algorithm. To increase the efficiency in optimising the variance of the Dirichlet prior, we relied on an initial hierarchy produced using Ward’s agglomerative clustering. The marginal likelihood of the root node was then used to select an appropriate scaling for the *β* hyperparameters. We subsequently refer to the combination of fast cluster initialization, Bayesian hierarchical clustering and prior hyperparameter optimisation using a symmetric or BAPS based prior as ‘fastbaps’.

### Conditioning on a pre-computed hierarchy

Given a precomputed hierarchy or phylogeny, we can use the recursion method described previously to decide when merging subclades of the tree is justified according to the DPM model. This allows us to identify a partition of the tree into clades that maximises the marginal likelihood of a DPM model given this hierarchy. As the hierarchy is pre-calculated, this approach is highly efficient and has a linear computational complexity in the number of nodes in the hierarchy. By leveraging the sparse matrix data structures in fastbaps and coupling them with the efficient implementation of Ward’s hierarchical clustering in the fastcluster package (24), we are able to generate clusterings of comparable quality to the full fastbaps algorithm very quickly (see Figure 4).

### Cluster stability using the Bootstrap

In order to determine the sensitivity of the resulting clusters to the alignment, we implemented a simple bootstrap procedure. For each bootstrap replicate, the loci are sampled with replacement and the fastbaps clustering algorithm is run. A binary similarity matrix is then produced as described in Strehl and Ghosh (25) where for each pair of sequences (i,j), the (i,j)th entry in the matrix is 1 if the ith and jth sequences are clustered together and 0 otherwise. As this matrix is invariant to label switching (26) and its dimension is invariant to the underlying number of clusters, the sum of the resulting matrices can be used to provide an indication of the stability of a given clustering of two isolates. This can be plotted in a heatmap alongside the dendrogram produced using the full algorithm to illustrate the robustness of the final clustering.

### Implementation

The fastbaps method is implemented using R and C++ and is available as an R package under the MIT open source licence at https://github.com/gtonkinhill/fastbaps. It can be run on Unix, Mac, or Windows operating systems and takes a multiple sequence alignment as input. The results can easily be parsed in R allowing for the generation of informative plots and further processing. The sparse matrix data structure generated by the package can also be used in other analysis types such as Principal Component Analysis (PCA) or k-means. All code to reproduce the simulations and results in this paper is available at https://github.com/gtonkinhill/fastbaps_manuscript.

### Simulation and evaluation

To compare the results of different algorithms, population structure was simulated using scrm as part of the Coala R package (27, 28). The number of underlying populations, recombination rate and migration rate were varied as described in Supplementary Table 1. Each parameter combination was run for 3 replicates providing 90 test datasets for analysis. A more detailed description of how the simulations were run can be found in the supplementary R notebook. Three clustering algorithms were considered in addition to the fastbaps approach. Snapclust and hierBAPS are available as R packages and were compared with fastbaps along with the fully Bayesian Structure algorithm (5, 13, 14, 29). Snapclust was run in parallel with the number of underlying clusters, *K*, varied between 2 and 30. The best fitting model was then chosen using either the Bayesian Information Criterion (BIC) or the Akaike Information Criterion (AIC). HierBAPS was run with 50 initial clusters. Finally, Structure was run with the no admixture model with 100,000 burn in iterations and 500,000 sampled iterations with 3 separate starting conditions. The run with highest likelihood was kept. The underlying number of clusters was fixed to the simulated *K* when running Structure to obtain a result in a reasonable amount of time, which nevertheless gives the method an unrealistic advantage against the alternatives which are fast enough to identify *K*.

To compare the simulated populations with those inferred by the different clustering algorithms, we used the Fowlkes-Mallows index (30). This accounts for both false positives and false negatives in determining the similarity between two clusterings and is defined as

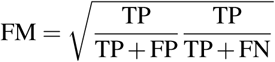

Here *T P* is the number of pairs of individuals that appear together in both clusterings, *F P* is the number that appear together in the inferred clusters but not the simulated and *F N* are those pairs that originated from the same population in the simulated data but are separated in the inferred clusters. Thus a clustering that perfectly matches the simulation with receive a Fowlkes-Mallows index of 1.

To compare with real data, six bacterial and two viral datasets were considered. The sets were chosen to cover a diverse range of pathogen species. The bacterial sets included the enteric bacteria *Escherichia coli*, the gram negative respiratory pathogen *Haemophilus influenzae*, as well as the firmicutes *Streptococcus pneumoniae*, *Listeria monocytogenes* and *Staphylococcus aureus* (31–35). The two viral datasets comprised a subset of sequences from the recent Ebola outbreak as well as a global dataset of over 110,000 HIV partial pol genes (positions 1-1497) obtained from the Los Alamos HIV Database (36, 37), with associated metadata on subtype and country of sampling. A summary of the size of each dataset is given in Supplementary Table 2. As there is no gold standard truth set for the real datasets we instead chose to compare the inferred clusters with a phylogeny built using Fasttree v2.1.10 (38). Here, we counted the number of pairs of isolates within each cluster that were inconsistent with the phylogeny in that there was an isolate belonging to another cluster within the clade representing their most recent common ancestor. These were considered ‘false positives’. This count was then divided by the total number of possible pairs within each cluster to give an indication of the error rate of the clustering (assuming the phylogeny to be correct). As the number of possible pairs increases with cluster size, this approach appropriately penalises clustering solutions with too many clusters.

## Results

### Fastbaps accurately clusters previously intractable viral and bacterial datasets containing thousands of samples

To investigate the performance of fastbaps on very large datasets, we analysed a dataset of over 110,000 HIV partial pol genes downloaded from the Los Alamos National Laboratories HIV Sequence Database (http://www.hiv.lanl.gov). The large number of sequences in this set presents a significant challenge to most model-based techniques as determining even the range of values of *K* is not straightforward. Additionally, we investigated the success of the algorithm on over 3, 100 pneumococcal genomes from the Maela refugee camp in Thailand (31). Whilst this dataset has fewer genomes, the large number of variable sites (284,194) makes this dataset very challenging for all current methods.

Figure 2 illustrates the resulting clusters inferred by fastbaps using the optimised BAPS prior on both the pneumococcal and HIV datasets. We made use of the UMAP dimensionality reduction method to project the isolates onto a 2D plane as this has been found to perform well in population genetics studies with very high numbers of individuals (39, 40). As the other model based-methods considered here were unable to produce results on either of these datasets in under a week, we compared the resulting clusters to those inferred using k-means. We used both the *K* identified by fastbaps as well as the *K* found using the ‘elbow method’ (41) when running the k-means algorithm. Figure 2a indicates that fastbaps mostly corresponds with the groupings formed in the UMAP projection. The visualization also demonstrates the utility of investigating the variability of the level of clonality across the clusters, as the layout of some clusters is more scattered spatially, reflecting the varying influence of recombination on the relatedness of pneumococcal strains. Whilst the k-means algorithm using the fastbaps inferred *K* (*K* = 79), provides a similar result, the *K* inferred using the elbow method leads to many inferred clusters being separated across the projection (Supplementary Figures 1 & 2). This highlights the advantages of using model-based techniques when investigating genetic datasets.

**Fig. 2.**
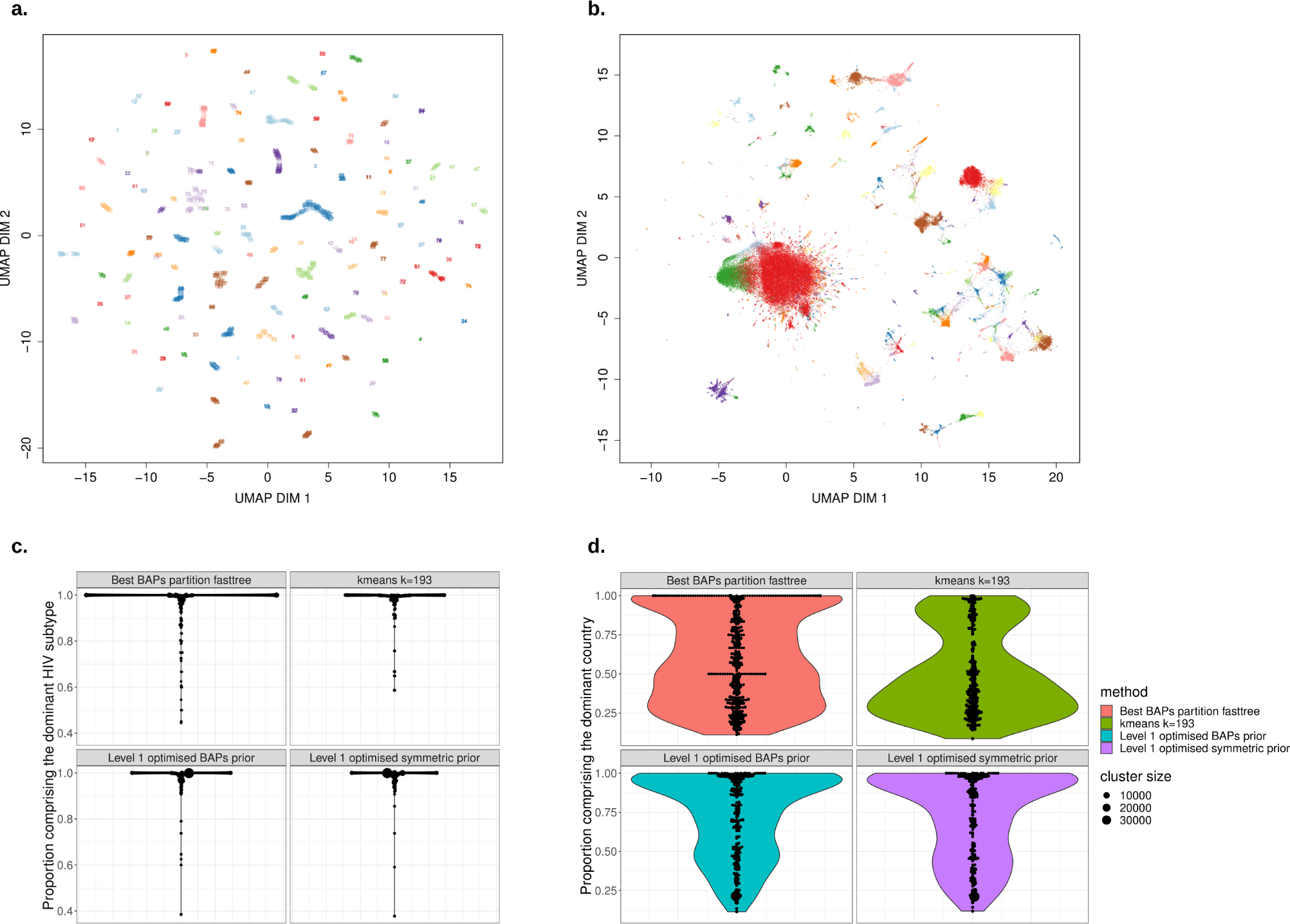
**a** A UMAP projection of over 3,100 pneumococcal genomes from Maela Thailand. The points are labeled and coloured by the cluster inferred using fastbaps with the optimised BAPS prior. In general the labeling appears to agree with the grouping observed in the UMAP projection. **b** Over 110,000 HIV-1 pol gene sequences from the Los Alamos public database. Sequences are coloured by their fastbaps inferred cluster. The large red cluster in the center contrasts with the noisy clustering observed using k-means (Supplementary Figure 3). The same plot with points numbered with their corresponding cluster is shown in Supplementary Figure 4. **c** Violin plots indicating the proportion of each inferred cluster that make up its most dominant subtype. Cluster size is represented by the size of a point and the width of the violin plot is proportional to the total number of clusters inferred. The fastbaps algorithm with the optimised BAPS prior provides a clustering that is most consistent with the underlying HIV subtypes whilst not overly segmenting the dataset. **d** Violin plots indicating the proportion of each inferred cluster that make up its most dominant country. Interestingly, the fastbaps approach is able to provide clusters that are more consistent with the underlying geography than alternative approaches such as k-means.

The fastbaps algorithm provides a cleaner clustering of the HIV dataset as seen in Figure 2b. Here, the UMAP dimensionality reduction technique as implemented in the umap-learn python package was used to project the isolates onto two dimensions (39). The large group of isolates in the centre of the UMAP projection has been assigned to one cluster. Conversely, k-means is unable to distinguish this group, and instead partitions it into many smaller overlapping clusters (Supplementary Figure 3 & 4). Similar to the pneumococcal dataset, when the lower value of *K* identified using the elbow method is used (*K* = 21), the resulting clusters are separated out over the UMAP projection (Supplementary Figure 5).

As HIV is highly recombinant, comparing the clustering of such a large global dataset to a phylogeny is unlikely to be informative. Instead, to further investigate the quality of the fastbaps clustering we examined the HIV subtype composition (inferred using a variety of methods, and including circulating recombinant forms as well as non-recombinant subtypes) as well as the distribution of the countries of origin of the isolates found in each cluster. Figure 2 indicates the proportion of each cluster that was dominated by a single subtype or any of its recombinant forms. Thus a cluster that is composed entirely of one subtype will have a proportion of 1. The figure indicates that despite having the same number of clusters as k-means, the fastbaps algorithm with both the optimised BAPS and optimised symmetric prior identifies larger clusters of purely one subtype.

This suggests the algorithm is identifying legitimate clusters within the dataset without over-partitioning the data. A similar comparison was made to investigate the proportion of each cluster that was dominated by a single country. Figure 2d indicates that fastbaps provides clusters that are more consistent with the underlying geography where as the k-means is dominated by many clusters that comprise isolates from many countries. Fastbaps was also used to partition a phylogeny. The resulting clustering produced a larger number of clusters due to the restriction of the phylogeny but the resulting clusters were more consistent by subtype and geography than those inferred using k-means.

### Simulations show that fastbaps cluster accuracy is the same as previous methods

In order to compare with different methods we made use of simulations where the true clustering solution was known. The Structure, hierBAPS, snapclust and fastbaps algorithms all achieved similar levels of accuracy on the simulated datasets (Figure 3). As expected, an increase in the migration rate between the simulated demes led to a subsequent decrease in the ability of the methods to accurately identify the underlying population structure. Additionally, a lower recombination rate led to poorer results, which is probably due to the algorithms correctly identifying additional phylogenetic structure within each cluster. Snapclust using AIC to select *K* as well as the fastbaps algorithm with the optimised BAPS prior achieved the best results over the simulated datasets. hierBAPS achieved a similar level of accuracy to the fastbaps algorithms using both the original (unscaled) BAPS prior as well as the optimised symmetric prior. Structure and Snapclust with BIC performed worst over the simulations. This was despite the MCMC chains of each Structure run showing good convergence characteristics (Supplementary Figure 6). A closer investigation of the Structure results indicated that it tended to allocate empty clusters despite being given the correct *K*. This suggests that both Snapclust with BIC and Structure tend to be overly conservative in estimating the number of underlying clusters *K*.

**Fig. 3.**
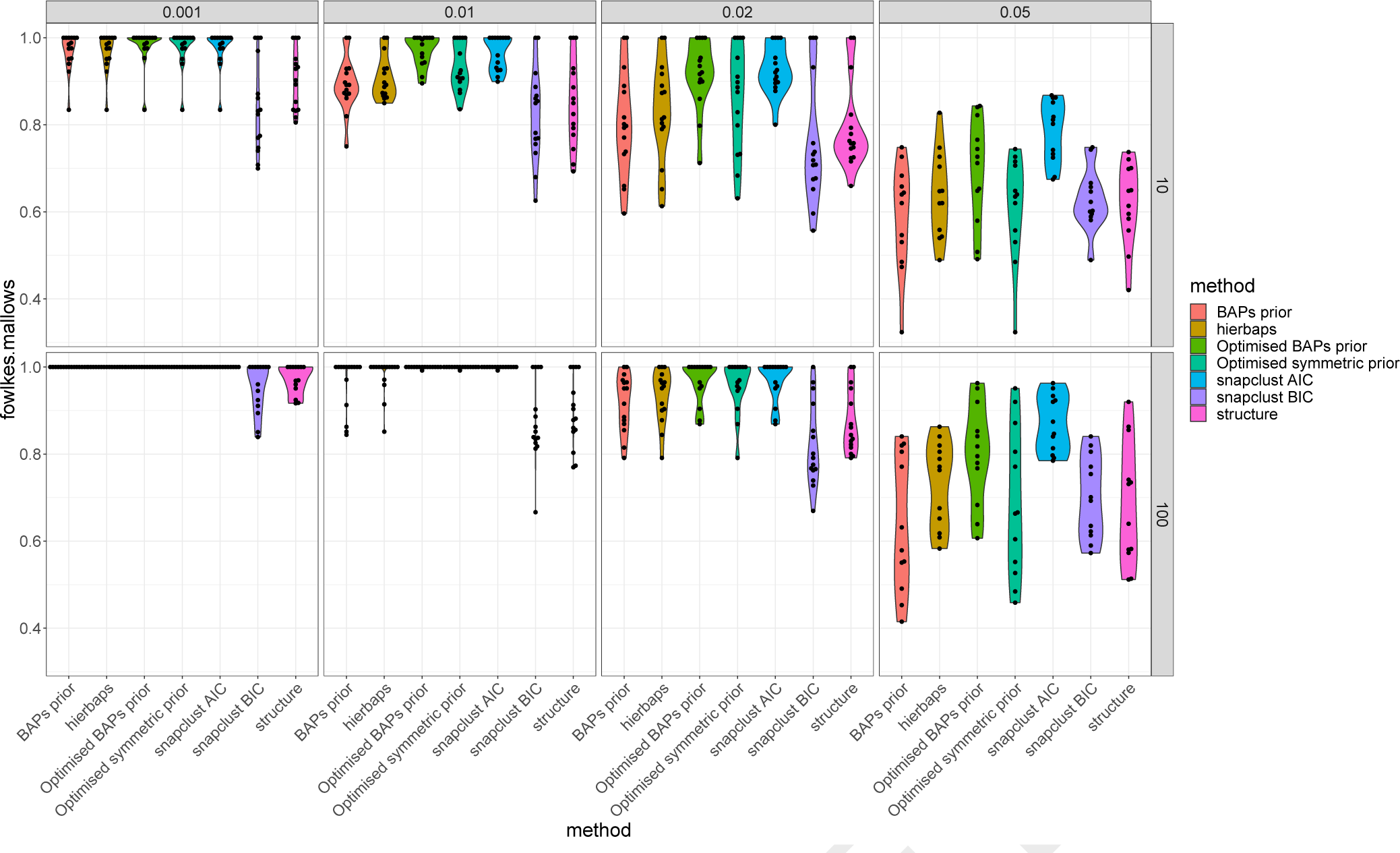
Violin plot indicating the Fowlkes-Mallows distance between the simulated clusters and those inferred by the different algorithms. Results for two distinct recombination rates are shown (10 and 100) as well as four different migration rates. The underlying number of clusters varied between 5-25. The plot indicates that as the migration rate increases the accuracy of the algorithm falls. Additionally, a lower recombination rate leads to lower accuracy, which is likely due to the algorithms identifying additional population structure within each simulated deme.

### Fastbaps is efficient and accurate on a diverse range of viral and bacterial datasets

To investigate the ability of each algorithm to detect population structure in real datasets, we made use of six additional bacterial and viral datasets of varying sizes and complexity (Table 2). Figure 4 indicates the resulting clusters inferred by the most promising algorithms from the simulation analysis on both a *Neisseria meningitidis* and *Haemophilus influenzae* data set. Here, we also provide comparisons with the prior from the original BHC algorithm and a partition of a Fasttree phylogeny using fastbaps. On the *H. influenzae* dataset, fastbaps using the optimised BAPS prior provided a clustering that was the most consistent with the phylogeny. Both the hierBAPS and snapclust solutions included polyphyletic clusters whilst the Bayesian Hierarchical Clustering (BHC) population mean based prior gave a similar result to the optimised BAPS solution. It should be noted that the initial fast clustering step was the same for both the BHC and optimised BAPS clusterings. The BHC prior failed to perform adequately on the large *N. meningitidis* dataset. Here, the BHC prior led to a highly partitioned solution with over 80 very small clusters. Conversely, the optimised BAPS prior led to a solution of similar quality to both hierBAPS and snapclust. Uncertainty in the Fasttree phylogenies was considered by comparing them with a bootstrapped phylogeny built using IQTREE v1.6.5 with the results found to be very similar. In both of these datasets the partition of the phylogeny using fastbaps provided a solution similar to that found using the optimised BAPS prior indicating it is an appropriate choice if a user’s goal is to simply partition a pre-calculated phylogeny.

**Fig. 4.**
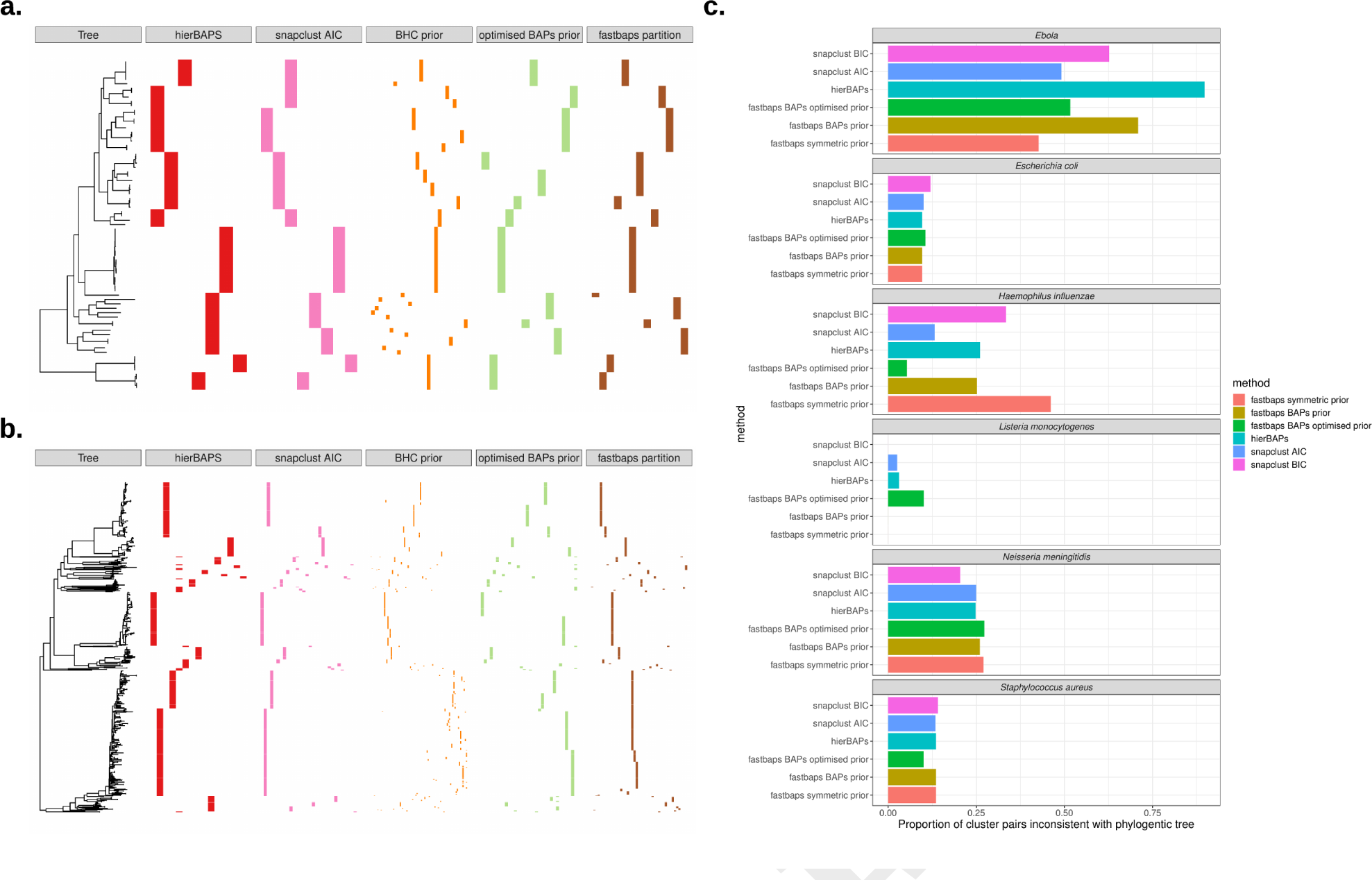
**a,b**. The resulting clustering inferred by hierBAPS, snapclust and fastbaps with both the original BHC prior and the optimised BAPS prior on both Staphylococcus aureus (**a**) and Neisseria meningitidis (**b**). The clusters are shown in comparison to a phylogeny generated using Fasttree. The final clustering indicates a partition of the Fasttree phylogeny using the fastbaps algorithm and thus is constrained to be consistent with the phylogeny. **c.** The proportion of isolate pairs that appear in the same inferred cluster but have isolates from a separate cluster in the clade represented by their most recent common ancestor in the Fasttree phylogeny. This provides a indication of the error of each algorithm assuming the phylogeny generated by Fasttree is correct. The fastbaps algorithm using either the optimised BAPS or symmetric prior outperforms the other methods on 5 out of 6 datasets.

Figure 4c, summarises the results of running the algorithms on all these data sets except the large HIV and pneumococcal collections. The figure indicates the number of pairs of isolates that are clustered together using the population structure algorithms but that are inconsistent with the phylogeny. Assuming the Fasttree phylogeny is accurate, this provides a measurement of error for each of the runs. The fastbaps algorithm with either the optimised symmetric or optimised BAPS priors archives a superior clustering on five out of six of the datasets. On the *N. meningitidis* dataset the snapclust algorithm using the BIC model selection criteria outperforms the other methods. The BIC model selection technique more heavily penalises solutions with a larger number of clusters and was in the simulations shown to frequently result in too few clusters. It is unlikely that the other methods are over-partitioning these data relative to snapclust with BIC. Hence, the outlying result for *N. meningitidis* could rather be due to inaccuracies in the Fasttree phylogeny, since *N. meningitidis* has a high recombination rate.

The computational run times and maximum memory requirements of each of the algorithms are shown in Tables 1 & 2 respectively. The tables indicate that fastbaps often performs more than an order of magnitude faster than snapclust and hierBAPS which themselves significantly outperform Structure in computational efficiency (13, 42). Part of this speed is due to the use of highly optimised sparse matrix libraries in fastbaps. As snapclust must be run for all values of *K* that are to be considered, its computational performance is directly tied to the maximum *K* chosen. Here, snapclust was run for a maximum *K* = 50 as this corresponded to a sensible upper bound for the *E. coli* dataset. Reducing the maximum *K* to use would improve the timing of the snapclust algorithm at the expense of failing to explore higher dimension solutions. However, Table 1 indicates that the fastbaps algorithm achieves a superior runtime compared with snapclust even for a single value of *K*, whilst automatically determining an appropriate number of clusters. Depending on the dataset, the prior optimisation step in fastbaps can take longer than running the complete algorithm. Thus if a short run-time is a necessity, running the method with a fixed prior is available as an option. Finally, if a hierarchy is already available, either through the use of a traditional phylogeny reconstruction algorithm such as Fasttree or via agglomerative clustering a partition using the fastbaps algorithm can be achieved with a complexity linear in the number of sequences. Fastbaps was able to partition a phylogeny of the full HIV dataset built using Fasttree (38) in 5.5 minutes.

**Table 1.**
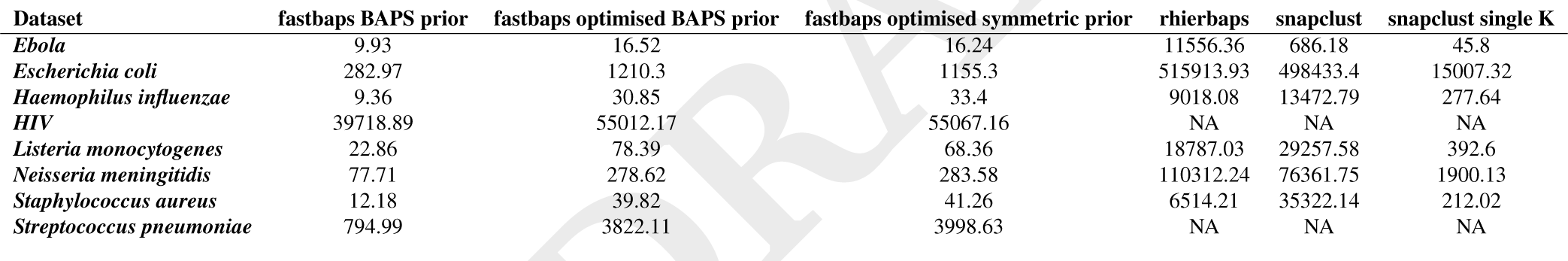
Total CPU time (seconds) of the different algorithms. Only fastbaps was able to run in under a week for the Pneumococcal and HIV datasets.

**Table 2.** Maximum memory usage of the different algorithms (Mb)

| Dataset | fastbaps BAPS prior | fastbaps optimised BAPS prior | fastbaps optimised symmetric prior | rhierBAPS | snapclust | snapclust single K |
| --- | --- | --- | --- | --- | --- | --- |
| *Ebola* | 411.64 | 468.64 | 487.32 | 1102.51 | 519.71 | 498.44 |
| *Escherichia coli* | 4290.98 | 4559.68 | 5178.42 | 15194.38 | 54714.39 | 45680.39 |
| *Haemophilus influenzae* | 394.56 | 375.84 | 382.09 | 852.5 | 4148.99 | 2113.79 |
| *HIV* | 3465.45 | 6511.35 | 7409.47 | NA | NA | NA |
| *Listeria monocytogenes* | 791.76 | 842.88 | 823.48 | 1400.69 | 5814.35 | 5194.14 |
| *Neisseria meningitidis* | 1439.4 | 1477.06 | 1382.16 | 4231.05 | 13779.11 | 10822.19 |
| *Staphylococcus aureus* | 384.85 | 387.04 | 396.44 | 742 | 3152.37 | 2423.9 |
| *Streptococcus pneumoniae* | 3915.3 | 5790.93 | 5710.96 | NA | NA | NA |

## Discussion

Finding clusters of related sequences present in a genetic alignment is a critical first step for many genetic and ecological analyses, allowing targeted sub-analysis and determination of the structure of the population when external classifications are not present. Model-based clustering methods are attractive due to their ability to hierarchically find highly specific clusters, estimate meaningful clustering parameters incorporating uncertainties, compare different model fits and produce probabilities of cluster assignment. Typical cohort sizes now range between thousands and tens of thousands of samples, growing even larger if combined with previous cohorts. However, the ability of methods to fit a clustering model to these alignments has not increased in step with these increases in alignment size (13, 29), with even modestly sized datasets requiring gene-by-gene or distance based approaches to determine clusters (4, 43, 44). Model-based approaches have also proved impractical in surveillance settings, where the continuous addition of small numbers of new sequences would require refitting the entire model.

By leveraging ideas from both the Bayesian Hierarchical Clustering (19) and the hierBAPS (14) algorithms, along with a fast sparse matrix based implementation, fastbaps enables efficient model-based clustering of large alignments that were previously infeasible to analyse using existing methods. Our approach enjoys all of the advantages inherent in modelbased approaches and can produce comparable or higher quality clusters than previous methods. Additionally, our algorithm is able to rapidly partition pre-computed phylogenies. This provides an attractive alternative approach for clustering genetic data when a phylogeny is available. The significant acceleration of inference provided by fastbaps enables the use of bootstrap replicates, allowing for the sensitivity of the resulting clusters to the alignment to be investigated in manner that has traditionally not been possible for model-based clustering methods on large alignments.

We verified our new approach by comparing its performance with other algorithms on both simulated data and eight real datasets. We show that as well as a considerable increase in the speed of the algorithm, we are able to achieve comparable accuracy, often outperforming previous model-based approaches. The speed of our new approach allowed us to cluster a large HIV sequence dataset containing over 100,000 sequences. The resulting clusters have higher concordance with HIV subtypes and geography than an alternative approach using k-means, whilst providing a principled method for selecting the underlying number of clusters; a major limitation of k-means.

Whilst our method enables the clustering of very large alignments, its complexity is still tied to the initial hierarchy generation and thus is *O*(*n*^2^) or possibly *O*(*nlog*(*n*)) if single linkage hierarchical clustering is used. After the initial hierarchy is generated, the remaining Bayesian hierarchical clustering is of the order *O*(*lm*^2^) where *m* is the number of clusters generated in the first stage and is the number of variable sites. Thus, in the future, as alignments begin to comprise tens of millions of sequences, additional improvements will need to be explored. Possible options include exploring the randomised extension to the BHC algorithm (45) or other machine learning based approaches such as those based on matrix decomposition (46).

Fastbaps therefore offers two new ways to rapidly find highquality clusters from a genetic alignment or phylogeny. The significant speed increase our software provides over previous approaches enables fitting a clustering model to previously intractable large alignments, and has the potential to allow continuous model refitting in surveillance settings. Fastbaps is provided as an open source package with a clearly documented interface.

## SOFTWARE AVAILABILITY

Source code available from: https://github.com/gtonkinhill/fastbaps Code for reproducing figures from:

https://github.com/gtonkinhill/fastbaps_manuscript Archived source code at time of publication to bioRxiv: https://doi.org/10.5281/zenodo.1472299

Archived code for figures at time of publication to bioRxiv: https://zenodo.org/badge/latestdoi/142294532

## ACKNOWLEDGEMENTS

This work was supported by the Wellcome Trust [206194] and [204016; to GTH; a Wellcome Trust PhD scholarship grant]; JC was supported by the ERC [742158]; and SDWF is supported in part by The Alan Turing Institute via an Engineering and Physical Sciences Research Council grant [EP/510129/1] and by the U.S. National Institutes of Health [R01AI135970].

## Supplementary Figures

**Supplementary Table S1:** Parameters used to simulate genetic population structure using scrm (27)

| Parameter | Value(s) |
| --- | --- |
| Number of replicates | 3 |
| Number of demes (clusters) | 5, 10, 15, 20, 25 |
| Deme level population size | 100 |
| Sequence length | 100,000 |
| Mutation rate | 0.01 |
| Recombination rate | 10, 100 |
| Migration rate | 0.001, 0.01, 0.02 |
| Proportion of population sampled | 0.1 |

**Supplementary Table S2:**
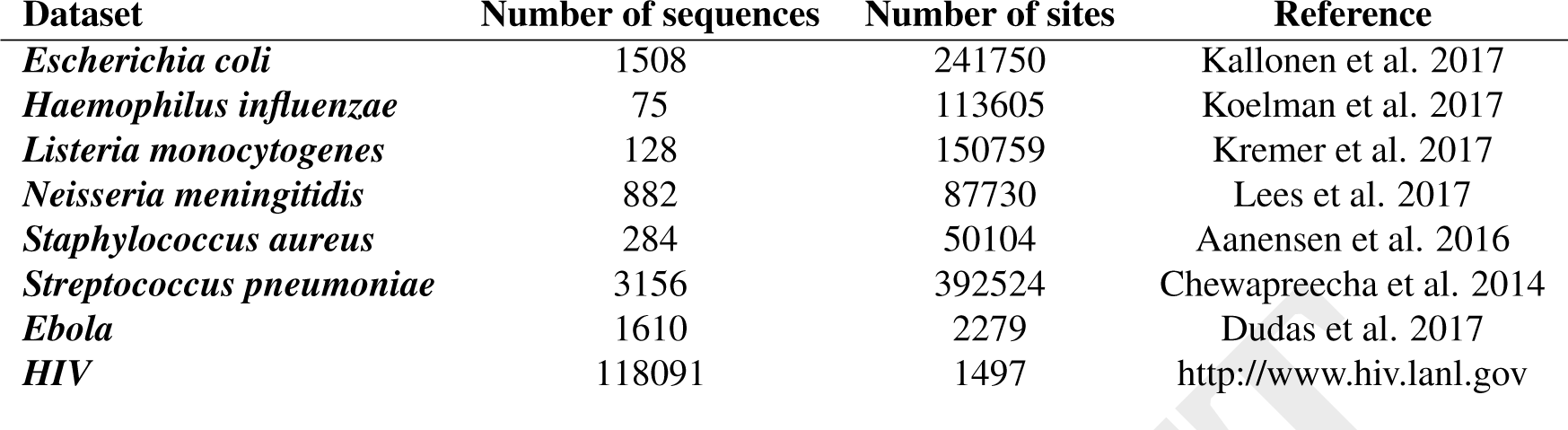
The datasets used in the comparison of different clustering methods. The HIV and pneumococcal datasets were only compared with k-means and fastbaps as they are too large for other model based methods.

| Dataset | Number of sequences | Number of sites | Reference |
| --- | --- | --- | --- |
| *Escherichia coli* | 1508 | 241750 | Kallonen et al. 2017 |
| *Haemophilus influenzae* | 75 | 113605 | Koelman et al. 2017 |
| *Listeria monocytogenes* | 128 | 150759 | Kremer et al. 2017 |
| *Neisseria meningitidis* | 882 | 87730 | Lees et al. 2017 |
| *Staphylococcus aureus* | 284 | 50104 | Aanensen et al. 2016 |
| *Streptococcus pneumoniae* | 3156 | 392524 | Chewapreecha et al. 2014 |
| *Ebola* | 1610 | 2279 | Dudas et al. 2017 |
| *HIV* | 118091 | 1497 | http://www.hiv.lanl.gov |

**Supplementary Figure 1:**
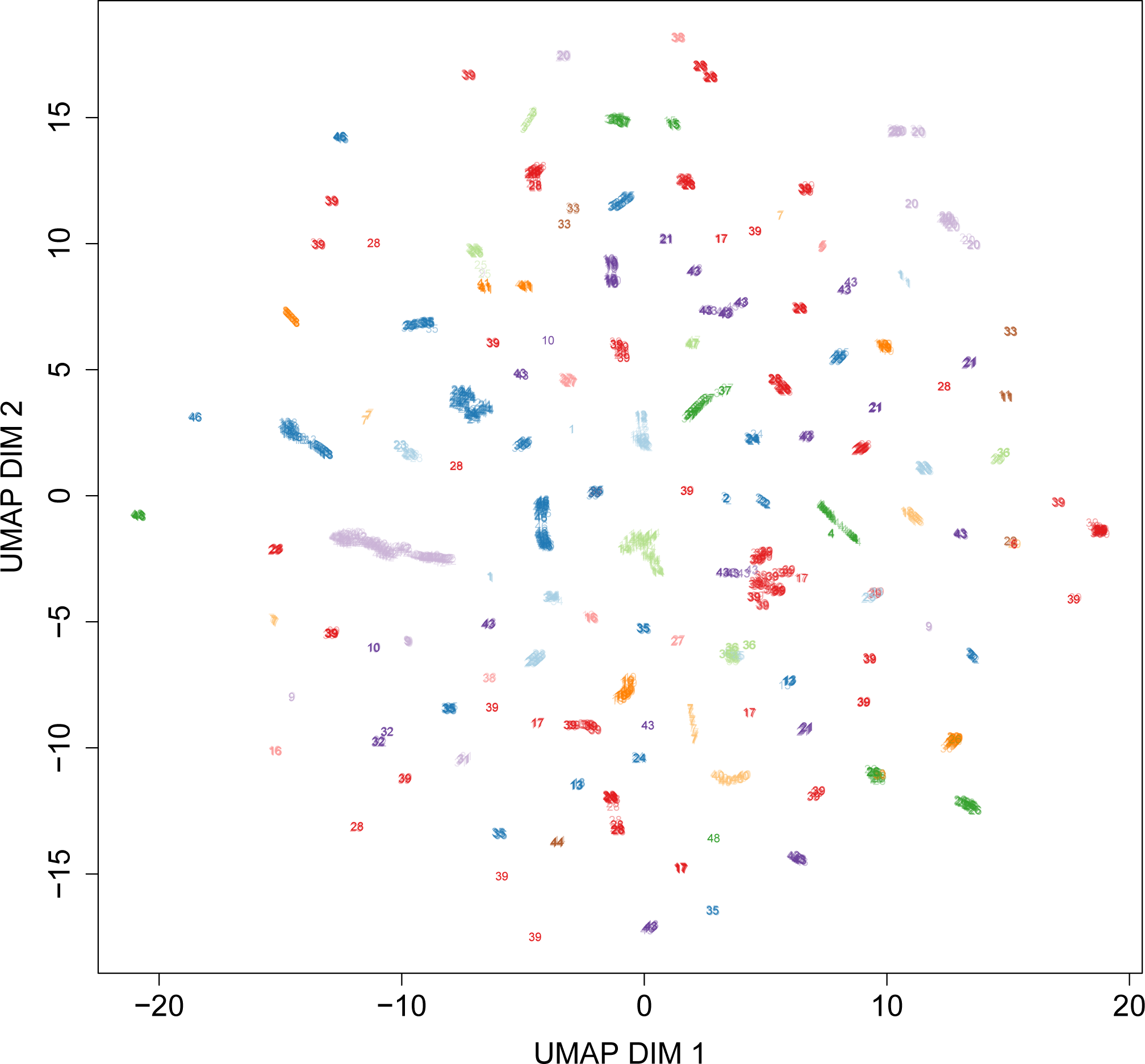
A UMAP plot of Pneumococcal isolates coloured and numbered by their inferred clustering using k-means with the number of underlying clusters found using the elbow method (*k* = 48).

**Supplementary Figure 2:**
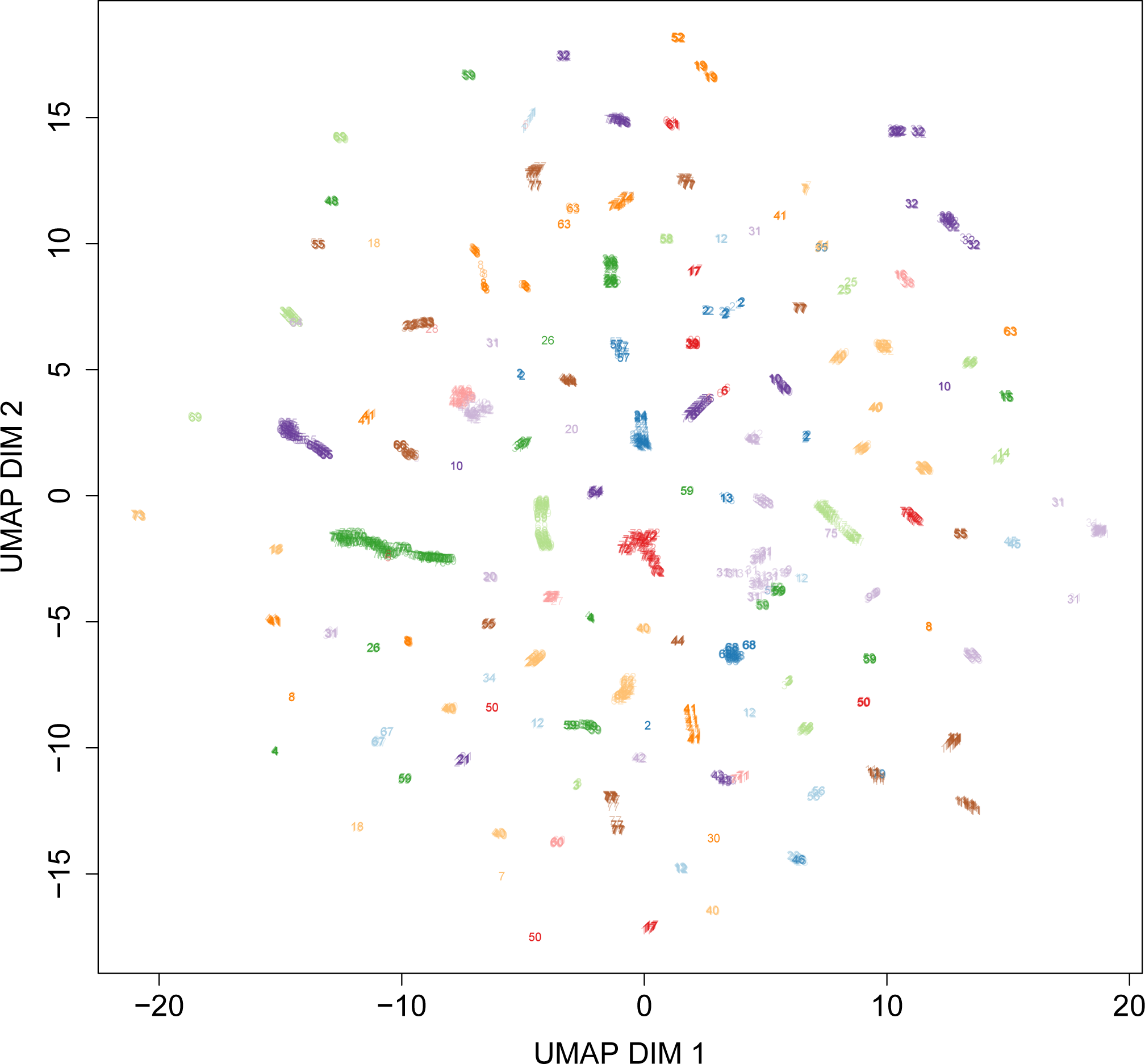
A UMAP plot of Pneumococcal isolates coloured and numbered by their inferred clustering using k-means with the number of underlying clusters matching that found using fastbaps (*k* = 79)

**Supplementary Figure 3:**
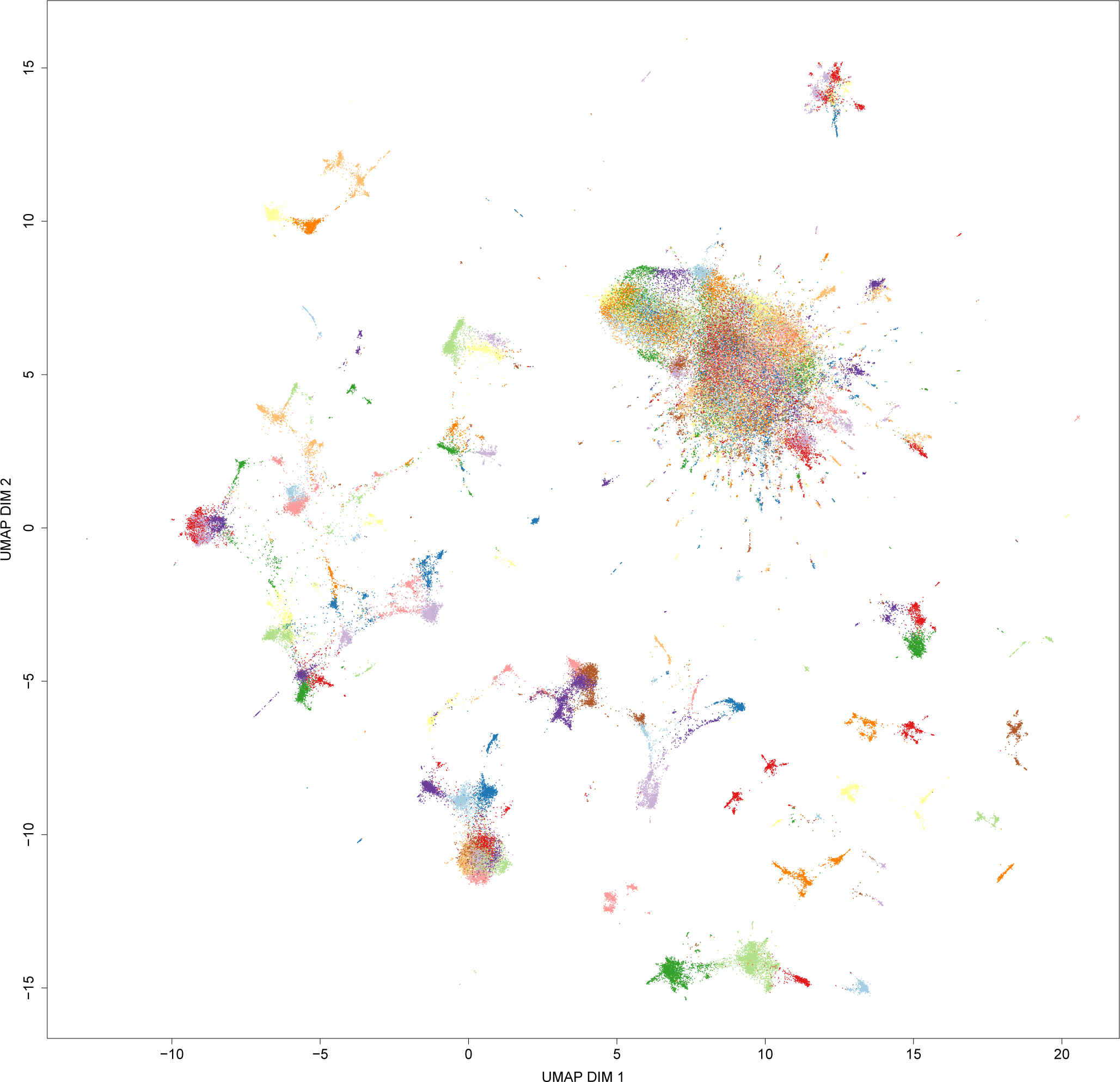
A UMAP plot of HIV isolates coloured by their inferred clustering using k-means with the number of underlying clusters matching that found using fastbaps (*k* = 193)

**Supplementary Figure 4:**
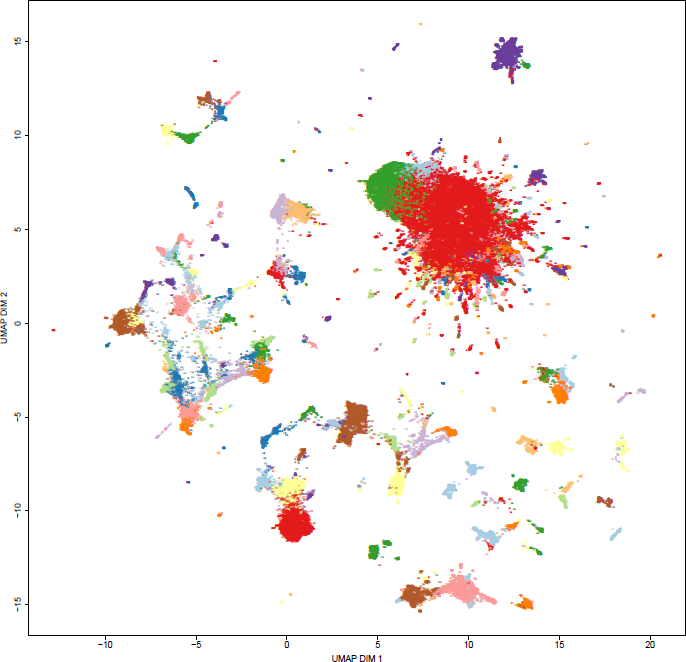
A UMAP plot of HIV isolates coloured and numbered by their inferred clustering using fastbaps with the optimised BAPS prior (*k* = 193)

**Supplementary Figure 5:**
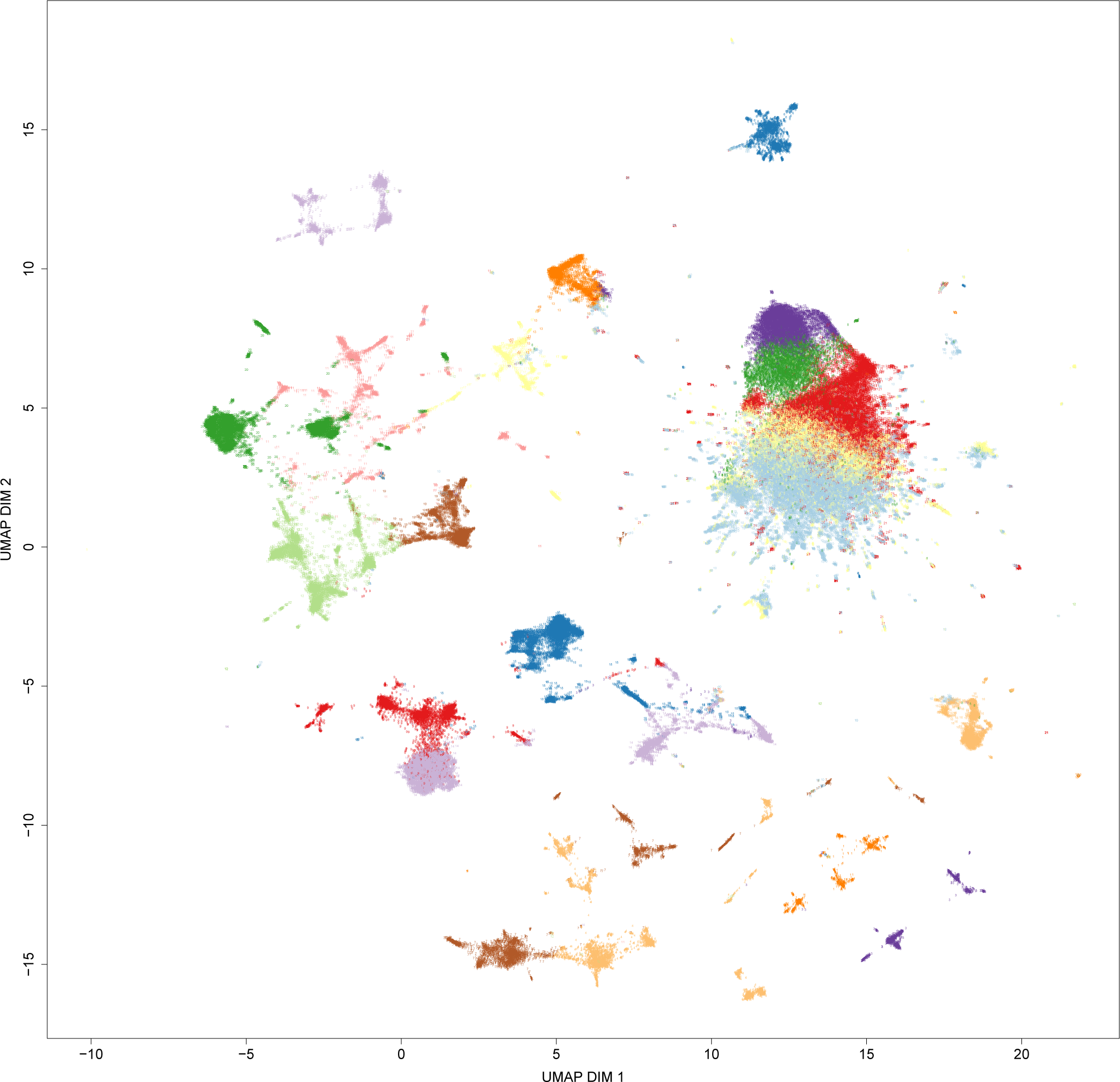
A UMAP plot of HIV isolates coloured and numbered by their inferred clustering using k-means with the number of underlying clusters found using the elbow method (*k* = 21)

**Supplementary Figure 6:**
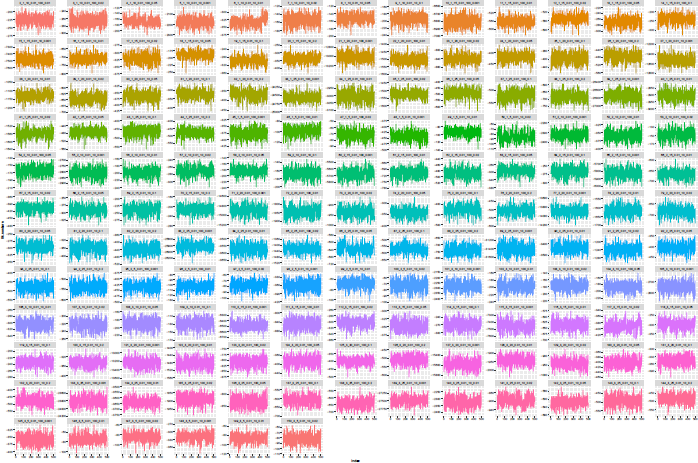
The MCMC chains for each run of the STRUCTURE algorithm on the simulated datasets

## Bibliography

1 Sergei L Kosakovsky Pond and Simon D W Frost. Not so different after all: a comparison of methods for detecting amino acid sites under selection. Mol. Biol. Evol., 22(5):1208–1222, May 2005.

2 John A Lees, Minna Vehkala, Niko Välimäki, Simon R Harris, Claire Chewapreecha, Nicholas J Croucher, Pekka Marttinen, Mark R Davies, Andrew C Steer, Steven Y C Tong, Antti Honkela, Julian Parkhill, Stephen D Bentley, and Jukka Corander. Sequence element enrichment analysis to determine the genetic basis of bacterial phenotypes. Nat. Commun., 7:12797, September 2016.

3 Sarah G Earle, Chieh-Hsi Wu, Jane Charlesworth, Nicole Stoesser, N Claire Gordon, Timothy M Walker, Chris C A Spencer, Zamin Iqbal, David A Clifton, Katie L Hopkins, Neil Woodford, E Grace Smith, Nazir Ismail, Martin J Llewelyn, Tim E Peto, Derrick W Crook, Gil McVean, A Sarah Walker, and Daniel J Wilson. Identifying lineage effects when controlling for population structure improves power in bacterial association studies. Nature Microbiology, (April):16041, April 2016.

4 Thibaut Jombart, Sébastien Devillard, and François Balloux. Discriminant analysis of principal components: a new method for the analysis of genetically structured populations. BMC Genet., 11:94, October 2010.

5 J K Pritchard, M Stephens, and P Donnelly. Inference of population structure using multilocus genotype data. Genetics, 155(2):945–959, June 2000.

6 Daniel Falush, Matthew Stephens, and Jonathan K Pritchard. Inference of population structure using multilocus genotype data: linked loci and correlated allele frequencies. Genetics, 164(4):1567–1587, August 2003.

7 Jukka Corander and Pekka Marttinen. Bayesian identification of admixture events using multilocus molecular markers. Mol. Ecol., 15(10):2833–2843, September 2006.

8 Jukka Corander, Pekka Marttinen, Jukka Sirén, and Jing Tang. Enhanced bayesian modelling in BAPS software for learning genetic structures of populations. BMC Bioinformatics, 9:539, December 2008.

9 Jukka Corander, Patrik Waldmann, and Mikko J Sillanpää. Bayesian analysis of genetic differentiation between populations. Genetics, 163(1):367–374, January 2003.

10 David H Alexander, John Novembre, and Kenneth Lange. Fast model-based estimation of ancestry in unrelated individuals. Genome Res., 19(9):1655–1664, September 2009.

11 Anil Raj, Matthew Stephens, and Jonathan K Pritchard. fastSTRUCTURE: variational inference of population structure in large SNP data sets. Genetics, 197(2):573–589, June 2014.

12 E C Anderson and E A Thompson. A model-based method for identifying species hybrids using multilocus genetic data. Genetics, 160(3):1217–1229, March 2002.

13 Marie-Pauline Beugin, Thibault Gayet, Dominique Pontier, Sébastien Devillard, and Thibaut Jombart. A fast likelihood solution to the genetic clustering problem. Methods Ecol. Evol., 9 (4):1006–1016, April 2018.

14 Lu Cheng, Thomas R Connor, Jukka Sirén, David M Aanensen, and Jukka Corander. Hierarchical and spatially explicit clustering of DNA sequences with BAPS software. Mol. Biol. Evol., 30(5):1224–1228, May 2013.

15 G Evanno, S Regnaut, and J Goudet. Detecting the number of clusters of individuals using the software STRUCTURE: a simulation study. Mol. Ecol., 14(8):2611–2620, July 2005.

16 H Akaike. A new look at the statistical model identification. IEEE Trans. Automat. Contr., 19 (6):716–723, December 1974.

17 Jerome Pella and Michele Masuda. The gibbs and split-merge sampler for population mixture analysis from genetic data with incomplete baselines. Can. J. Fish. Aquat. Sci., 63(3): 576–596, April 2011.

18 John P Huelsenbeck, Peter Andolfatto, and Edna T Huelsenbeck. Structurama: bayesian inference of population structure. Evol. Bioinform. Online, 7:55–59, June 2011.

19 Katherine A Heller and Zoubin Ghahramani. Bayesian hierarchical clustering. In Proceedings of the 22Nd International Conference on Machine Learning, ICML ‘05, pages 297–304, New York, NY, USA, 2005. ACM.

20 Anthony R Florita, Larry J Brackney, Todd P Otanicar, and Jeffrey Robertson. Classification of commercial building electrical demand profiles for energy storage applications. J. Sol. Energy Eng., 135(3):031020, August 2013.

21 Richard S Savage, Katherine Heller, Yang Xu, Zoubin Ghahramani, William M Truman, Murray Grant, Katherine J Denby, and David L Wild. R/BHC: fast bayesian hierarchical clustering for microarray data. BMC Bioinformatics, 10:242, August 2009.

22 Fionn Murtagh and Pierre Legendre. Ward’s hierarchical agglomerative clustering method: Which algorithms implement ward’s criterion? J. Classification, 31(3):274–295, October 2014.

23 Marek Gagolewski, Maciej Bartoszuk, and Anna Cena. Genie: A new, fast, and outlier-resistant hierarchical clustering algorithm. Inf. Sci., 363:8–23, October 2016.

24 Daniel Müllner and Others. fastcluster: Fast hierarchical, agglomerative clustering routines for R and python. J. Stat. Softw., 53(9):1–18, 2013.

25 Alexander Strehl and Joydeep Ghosh. Cluster ensembles — a knowledge reuse framework for combining multiple partitions. J. Mach. Learn. Res., 3(Dec):583–617, 2002.

26 Sylvia Richardson and Peter J Green. On bayesian analysis of mixtures with an unknown number of components (with discussion). Journal of the Royal Statistical Society: series B (statistical methodology), 59(4):731–792, 1997.

27 Paul R Staab, Sha Zhu, Dirk Metzler, and Gerton Lunter. scrm: efficiently simulating long sequences using the approximated coalescent with recombination. Bioinformatics, 31(10): 1680–1682, May 2015.

28 Paul R Staab and Dirk Metzler. Coala: an R framework for coalescent simulation. Bioinformatics, 32(12):1903–1904, June 2016.

29 Gerry Tonkin-Hill, John A Lees, Stephen D Bentley, Simon D W Frost, and Jukka Corander. RhierBAPS: An R implementation of the population clustering algorithm hierBAPS. Wellcome Open Res, 3:93, July 2018.

30 E B Fowlkes and C L Mallows. A method for comparing two hierarchical clusterings. J. Am. Stat. Assoc., 78(383):553–569, September 1983.

31 Claire Chewapreecha, Simon R Harris, Nicholas J Croucher, Claudia Turner, Pekka Marttinen, Lu Cheng, Alberto Pessia, David M Aanensen, Alison E Mather, Andrew J Page, Susannah J Salter, David Harris, Francois Nosten, David Goldblatt, Jukka Corander, Julian Parkhill, Paul Turner, and Stephen D Bentley. Dense genomic sampling identifies highways of pneumococcal recombination. Nat. Genet., 46(3):305–309, March 2014.

32 David M Aanensen, Edward J Feil, Matthew T G Holden, Janina Dordel, Corin A Yeats, Artemij Fedosejev, Richard Goater, Santiago Castillo-Ramírez, Jukka Corander, Caroline Colijn, Monika A Chlebowicz, Leo Schouls, Max Heck, Gerlinde Pluister, Raymond Ruimy, Gunnar Kahlmeter, Jenny Åhman, Erika Matuschek, Alexander W Friedrich, Julian Parkhill, Stephen D Bentley, Brian G Spratt, Hajo Grundmann, and European SRL Working Group. Whole-Genome sequencing for routine pathogen surveillance in public health: a population snapshot of invasive staphylococcus aureus in europe. MBio, 7(3), May 2016.

33 John A Lees, Philip H C Kremer, Ana S Manso, Nicholas J Croucher, Bart Ferwerda, Mercedes Valls Serón, Marco R Oggioni, Julian Parkhill, Matthijs C Brouwer, Arie van der Ende, Diederik van de Beek, and Stephen D Bentley. Large scale genomic analysis shows no evidence for pathogen adaptation between the blood and cerebrospinal fluid niches during bacterial meningitis. Microb Genom, 3(1):e000103, January 2017.

34 P H C Kremer, J A Lees, M M Koopmans, B Ferwerda, A W M Arends, M M Feller, K Schipper, M Valls Seron, A van der Ende, M C Brouwer, D van de Beek, and S D Bentley. Benzalkonium tolerance genes and outcome in listeria monocytogenes meningitis. Clin. Microbiol. Infect., 23(4):265.e1–265.e7, April 2017.

35 D Koelman, P Kremer, J Lees, M Brouwer, S Bentley, and D van de Beek. Bacterial hypervirulence in haemophilus influenzae meningitis identified by whole genome sequencing. J. Neurol. Sci., 381:181–182, October 2017.

36 Los Alamos National Laboratory. HIV databases. https://www.hiv.lanl.gov/content/index. Accessed: 2018-10-25.

37 Gytis Dudas, Luiz Max Carvalho, Trevor Bedford, Andrew J Tatem, Guy Baele, Nuno R Faria, Daniel J Park, Jason T Ladner, Armando Arias, Danny Asogun, Filip Bielejec, Sarah L Caddy, Matthew Cotten, Jonathan D’Ambrozio, Simon Dellicour, Antonino Di Caro, Joseph W Diclaro, Sophie Duraffour, Michael J Elmore, Lawrence S Fakoli, Ousmane Faye, Merle L Gilbert, Sahr M Gevao, Stephen Gire, Adrianne Gladden-Young, Andreas Gnirke, Augustine Goba, Donald S Grant, Bart L Haagmans, Julian A Hiscox, Umaru Jah, Jeffrey R Kugelman, Di Liu, Jia Lu, Christine M Malboeuf, Suzanne Mate, David A Matthews, Christian B Matranga, Luke W Meredith, James Qu, Joshua Quick, Suzan D Pas, My V T Phan, Georgios Pollakis, Chantal B Reusken, Mariano Sanchez-Lockhart, Stephen F Schaffner, John S Schieffelin, Rachel S Sealfon, Etienne Simon-Loriere, Saskia L Smits, Kilian Stoecker, Lucy Thorne, Ekaete Alice Tobin, Mohamed A Vandi, Simon J Watson, Kendra West, Shannon Whitmer, Michael R Wiley, Sarah M Winnicki, Shirlee Wohl, Roman Wölfel, Nathan L Yozwiak, Kristian G Andersen, Sylvia O Blyden, Fatorma Bolay, Miles W Carroll, Bernice Dahn, Boubacar Diallo, Pierre Formenty, Christophe Fraser, George F Gao, Robert F Garry, Ian Goodfellow, Stephan Günther, Christian T Happi, Edward C Holmes, Brima Kargbo, Sakoba Keïta, Paul Kellam, Marion P G Koopmans, Jens H Kuhn, Nicholas J Loman, N’faly Magassouba, Dhamari Naidoo, Stuart T Nichol, Tolbert Nyenswah, Gustavo Palacios, Oliver G Pybus, Pardis C Sabeti, Amadou Sall, Ute Ströher, Isatta Wurie, Marc A Suchard, Philippe Lemey, and Andrew Rambaut. Virus genomes reveal factors that spread and sustained the ebola epidemic. Nature, 544(7650):309–315, April 2017. ISSN 0028-0836, 1476-4687. doi: 10.1038/nature22040.

38 Morgan N Price, Paramvir S Dehal, and Adam P Arkin. FastTree 2 – approximately Maximum-Likelihood trees for large alignments. PLoS One, 5(3):e9490, March 2010.

39 Leland McInnes and John Healy. UMAP: Uniform manifold approximation and projection for dimension reduction. February 2018.

40 Alex Diaz-Papkovich, Luke Anderson-Trocme, and Simon Gravel. Revealing multi-scale population structure in large cohorts. September 2018.

41 Alboukadel Kassambara. Practical Guide to Cluster Analysis in R: Unsupervised Machine Learning. STHDA, August 2017.

42 Emily K Latch, Guha Dharmarajan, Jeffrey C Glaubitz, and Olin E Rhodes. Relative performance of bayesian clustering software for inferringpopulation substructure and individual assignment at low levels of population differentiation. Conserv. Genet., 7(2):295–302, April 2006.

43 Ji Zhang, Jani Halkilahti, Marja-Liisa Hänninen, and Mirko Rossi. Refinement of wholegenome multilocus sequence typing analysis by addressing gene paralogy. J. Clin. Microbiol., 53(5):1765–1767, May 2015.

44 John A Lees, Simon R Harris, Gerry Tonkin-Hill, Rebecca A Gladstone, Stephanie Lo, Jeffrey N Weiser, Jukka Corander, Stephen D Bentley, and Nicholas J Croucher. Fast and flexible bacterial genomic epidemiology with PopPUNK. July 2018.

45 Zoubin Ghahramani Katherine A. Heller. Randomized algorithms for fast bayesian hierarchical clustering. citeseerx.ist.psu.edu/viewdoc/summary?doi=10.1.1.60.298, 2005.

46 Zhirong Yang, Jukka Corander, and Erkki Oja. Low-Rank doubly stochastic matrix decomposition for cluster analysis. J. Mach. Learn. Res., 17(187):1–25, 2016.

